# Single-cell resolution uncovers cell type-specific dysregulation in Parkin-deficient neuron-microglia co-cultures

**DOI:** 10.64898/2026.03.27.714690

**Authors:** Evelyn Knappe, Kristian Händler, Linn Streubel-Gallasch, Franziska Rudolph, Daniel Alvarez Fischer, Sally A. Cowley, Anne Grünewald, Malte Spielmann, Christine Klein, Philip Seibler

**Affiliations:** Institute of Neurogenetics, University of Lübeck and University Hospital Schleswig-Holstein, Ratzeburger Allee 160, 23538, Lübeck, Germany; Institute of Human Genetics, University of Lübeck and University Hospital Schleswig-Holstein, Ratzeburger Allee 160, 23538, Lübeck, Germany; James and Lillian Martin Centre for Stem Cell Research, Sir William Dunn School of Pathology, University of Oxford, Oxford, United Kingdom; Luxembourg Centre for Systems Biomedicine, University of Luxembourg, Esch-sur-Alzette, Luxembourg; Max Planck Institute for Molecular Genetics, Berlin, Germany; Institute for Medical and Human Genetics, Charité – Universitätsmedizin Berlin, Berlin, Germany

**Keywords:** iPSC, dopaminergic neurons, microglia, co-culture, single-cell RNA sequencing, Parkin, *PRKN*, Parkinson’s disease

## Abstract

**Background:** Mutations in the E3 ubiquitin ligase Parkin (encoded by *PRKN*) are the most frequently known cause of recessively inherited Parkinson’s disease. In addition to the loss of dopaminergic neurons, microglial activation is another pathological feature observed in Parkinson’s disease. While postmortem brain samples show the end stage of the disease, neurons and glia derived from patients’ induced pluripotent stem cells (iPSCs) provide a model for detecting early pre-degenerative disease trajectories. However, mixed cell populations often confound these cultures, leading to heterogeneous disease phenotypes.

**Methods:** Here, we tease apart the cell type-specific phenotypes underlying Parkin-linked Parkinson’s disease by performing single-nucleus RNA sequencing in iPSC-derived co-cultures of dopaminergic neurons and microglia from *PRKN* mutation carriers and healthy controls. We validated our transcriptomic key findings through inflammatory cytokine profiling and live-cell calcium imaging.

**Results:** Single-nucleus RNA sequencing identified seven major cell types composed of neuronal, glial, and precursor cells, with dopaminergic neurons accounting for the largest cell population. Pathway analysis revealed cell type-specific dysregulated biological processes in Parkin-deficient cells, including gene expression differences in dopaminergic neurons that control mitophagy and dopamine homeostasis, whereas microglia showed changes in calcium homeostasis and inflammatory signaling. Functional analysis verified elevated secretion of monocyte chemotactic protein 1 in *PRKN*-mutant co-cultures compared with controls, linking Parkin deficiency to increased microglial chemotactic signaling. Furthermore, lower intracellular calcium levels and diminished calcium release following treatment confirmed impaired calcium homeostasis in *PRKN*-mutant microglia.

**Conclusions:** Profiling at the single-cell level resolved distinct cell subpopulations, enabling us to identify cell type-specific pathway disturbances underlying Parkin deficiency. This unique dataset provides a basis for understanding the impairment of individual cell types and the impact of cellular crosstalk in Parkinson’s disease pathology.

## BACKGROUND

Aside from the degeneration of dopaminergic (DA) neurons and the subsequent reduction in dopamine levels, microglial activation has been observed in patients with Parkinson’s disease (PD) and in post-mortem brain tissue [1–5]. However, the molecular mechanisms underlying the pathology of PD in DA neurons and microglia remain poorly understood. By the time a clinical diagnosis is confirmed, it is estimated that 40-60% of DA neurons have already degenerated [6], and several inflammatory markers are detectable at this stage [5]. Therefore, model systems, elucidating the disease’s pre-degenerative phase in biologically relevant cell types, are required to detect early and targetable disease trajectories to prevent neurodegeneration.

While postmortem brain samples provide insight into the end stage of the disease, induced pluripotent stem cell (iPSC) technology enables the study of patient-derived disease-relevant cell types before disease onset, retaining the entire genetic background. Mutations in the E3 ubiquitin ligase Parkin (encoded by *PRKN*) are the most frequently known cause of recessively inherited PD [7]. Work with iPSC-derived DA neurons from PD patients harboring mutations in *PRKN* revealed its role in mitochondrial function and biogenesis [8–12], mitophagy [13, 14], DA homeostasis [15, 16], calcium homeostasis [17], and inflammation [18]. However, iPSC-derived neuronal cultures are often confounded by a heterogeneous appearance of disease phenotypes *in vitro* due to a mixed population of cells with different degrees of maturation [19]. While bulk analyses across various cell types yield only a population average, profiling at the single-cell level can exploit population heterogeneity to resolve distinct cell subpopulations, offering a unique approach to understanding PD trajectories.

To gain a better understanding of how Parkin deficiency impacts pathways in individual cell types, we performed single-nucleus profiling of *PRKN*-mutant and control iPSC-derived DA neuron-microglia co-cultures.

## METHODS

### Patients and iPSC lines

iPSC lines were previously derived from three healthy individuals (Table S1) and three patients with biallelic *PRKN*-linked PD (Table S2). Control lines are registered in hPSCreg, with accompanying quality control reports, and patient lines were characterized in this study (Figure S1). iPSCs were plated onto Matrigel (Corning Inc, Glendale, USA)-coated plates and cultured in mTeSR1 medium (STEMCELL Technologies, Vancouver, Canada). Cells were passaged using 0.5 mM EDTA in PBS every 4-5 days. Whole-genome single-nucleotide polymorphism (SNP) analysis was performed for iPSC lines and parental fibroblasts on the Infinium OmniExpress-24-Bead Chip (Illumina, San Diego, USA) and analyzed with the software KaryoStudio (Illumina). Karyotypic aberrations were not detected (data not shown).

### Differentiation into dopaminergic neurons

The differentiation into DA neurons was performed according to a previously established protocol [20], with slight modifications [21]. In brief, neuronal differentiation of iPSCs was initiated by adding the SMAD pathway inhibitors SB431542 (10 mM, Tocris, Bristol, UK) and LDN193189 (100 nM, Stemgent, Cambridge, USA). Dopaminergic patterning was induced by adding recombinant sonic hedgehog (100 ng/ml, STEMCELL Technologies), purmorphamine (2 µM, STEMCELL Technologies), fibroblast growth factor 8a (200 ng/ml, STEMCELL Technologies), and CHIR99021 (3 µM, STEMCELL Technologies). From day 13 of differentiation, the medium contained brain-derived neurotrophic factor (20 ng/ml, STEMCELL Technologies), ascorbic acid (0.2 mM, Sigma Aldrich, St. Louis, USA), glial cell line-derived neurotrophic factor (20 ng/ml, STEMCELL Technologies), transforming growth factor β3 (1 ng/ml, Peprotech, Cranbury, USA), cyclic adenosine monophosphate (cAMP; 0.5 mM, Enzo Lifesciences, Farmingdale, USA), and DAPT (10 µM, Tocris). At day 21, 3.5*10E5 cells were plated in high-density drops onto dishes coated with poly-D-lysine/laminin. A medium change was performed every 2-3 days. Neuronal differentiation factors were withdrawn on day 40 of differentiation. This also includes cAMP, which has been shown to be toxic for microglia [22].

### Differentiation into neuron-microglia co-cultures

Differentiation of iPSCs into microglial precursors was performed as described previously [23, 24], with minor modifications [21]. In brief, iPSCs were singled and seeded into a 24-well Aggrewell 800 plate (STEMCELL Technologies) at a density of 4*10E6 cells/well. For mesodermal induction, embryoid body (EB) formation was complemented by supplying bone morphogenetic protein 4 (50 ng/ml, Peprotech), stem cell factor (20 ng/ml, Peprotech), and vascular endothelial growth factor (50 ng/ml, Peprotech). After the transfer of the EBs into T75 cell culture flasks, myeloid progenitor cells were formed with the support of Interleukin-3 (IL-3) (25 ng/ml, STEMCELL Technologies) and macrophage colony-stimulating factor (100 ng/ml, STEMCELL Technologies) and released into the medium as suspension cells. The medium was changed once a week. At day 30 of differentiation, myeloid progenitor cells were collected and plated onto 50-day-old DA neurons with a density of 4*10E5 cells/24-well. The maturation medium was supplemented with 100 ng/ml of IL-34. Half medium changes were performed every 48h. After 14 days, the mature co-cultures were used for further analysis.

### Western blot analysis

Protein extraction and western blot analysis were performed according to previously published protocols [25]. In brief, cells were lysed in RIPA buffer (25 mM TRIS-HCl pH 7.6, 150 mM NaCl, 1% NP-40, 0.1% SDS). Protein concentration was measured using the DC Protein Assay (Biorad, Hercules, USA) according to the manufacturer’s instructions. For western blot analysis, 10 µg of protein was loaded on NuPAGE™ 4-12% Bis-Tris gels (Thermo Fisher Scientific, Waltham, USA) and transferred onto a nitrocellulose membrane. Blocking was performed for 1h at room temperature (RT) in 1% milk in TBST (TBS, 1% Tween-20). The membrane was incubated overnight at 4°C with primary antibodies diluted in 1% milk. Secondary antibodies (Thermo Fisher Scientific) were diluted 1:8000 in 1% milk and membranes incubated for 1h at RT. All primary antibodies are listed in Table S3. Proteins were visualized using Supersignal West Pico Luminescent Substrate (Thermo Fisher Scientific) according to the manufacturer’s instructions.

### Cytokine release measurements

Co-cultures were treated with different stressors 10 ng/ml IL-1β (Peprotech), 50 ng/ml tumor necrosis factor alpha (TNF-α, Peprotech), or 100 ng/ml of interferon gamma (IFN-γ, Peprotech) and LPS (Sigma-Aldrich, St. Louis, USA) for 24h at 37°C to induce cytokine release as described previously [21]. To determine basal levels of cytokines, we measured untreated and undiluted samples. All media samples were stored at −80°C until analysis. The Legendplex Human Inflammation Panel 1 assay (Biolegend, San Diego, USA) was used to assess the concentration of 13 pro- and anti-inflammatory cytokines simultaneously in the cell culture media samples according to the manufacturer’s instructions. All measurements were performed in duplicates on the spectral flow cytometer Cytek Aurora (Cytek Biosciences, Fremont, USA). IL-33 was not detected and excluded from the analysis. An online tool provided by the manufacturer (https://legendplex.qognit.com (assessed on 24 April 2023)) was used to analyze all data. Following gating, the individual sample data were matched to the standard curves for the respective cytokines, enabling direct quantification of cytokine content for each sample. Subsequently, cytokine concentrations were normalized to the respective protein concentration of each sample.

### Immunofluorescence staining

Cells were fixed with 4% paraformaldehyde for 10 min at RT [25] and subsequently permeabilized and blocked with PBS containing 4% normal donkey (Sigma-Aldrich) or goat serum (Thermo Fisher Scientific), 0.1% bovine serum albumin (BSA), 0.1% Triton X-100, and 0.05% sodium azide for 1h at RT. Cells were incubated with primary antibodies (Table S3) overnight at 4°C. Incubation with secondary antibodies was performed in PBS with 3% BSA and 0.05% sodium azide for 1h at RT. DAPI Fluoromount-G (Southern Biotech, Birmingham, USA) was used as a mounting medium. Images were taken using the confocal laser scanning microscope LSM 900 (Zeiss, Jena, Germany).

### Single-nucleus RNA-seq

To profile the gene expression of co-cultures, single-nucleus transcriptome analysis was performed on all cell lines. In brief, 14-day-old co-cultures were pelleted and stored at −80°C. Nuclei isolation was performed using the Chromium Nuclei Isolation Kit with RNase Inhibitor (10x Genomics, Pleasanton, USA) according to manufacturer’s recommendations. Subsequently, nuclei were fixed by using Chromium Next GEM Single Cell Fixed RNA Sample preparation Kit supplemented with 4% formaldehyde. To isolate intact nuclei from cell debris and ambient RNA, the fixed single nuclei solution was sorted using a BD FACSAria III system and DAPI staining.

RNA gene expression libraries were generated using the Chromium Fixed RNA Kit chemistry with the Human Transcriptome probe set according to the manufacturer’s recommendations. Individual samples were multiplexed into 2 libraries of 3 samples, each using different probe set barcodes, followed by nuclei counting and mixing equal numbers from different replicates. A 10x Chromium X device was used for microfluidic single nuclei compartmentalization via Gel bead-in-emulsion (GEM) creation with subsequent library prep according to the manufacturer’s instructions. The two resulting libraries were quantified by qPCR using the NEBNext Library Quant Kit for Illumina (New England Biolabs, Ipswich, USA) on a 7300 Real Time PCR system (Applied Biosystems) and library size distribution was determined by Agilent High Sensitivity DNA assay on a Bioanalyzer 2100 device (Agilent, Santa Clara, USA). The libraries were equimolarly pooled and sequenced using NextSeq 2000 P3 Reagents (100 Cycles) on a NextSeq 2000 device (Illumina). Data was converted to FASTQ format, and libraries were demultiplexed using bcl2fastq2 v2.20 (Illumina). Subsequently, the data was further demultiplexed into individual samples and converted into single cell count matrices using cellranger 7.0.0 (10x Genomics) in conjunction with the human reference transcriptome GRCh38-2020-A and Chromium_Human_Transcriptome_Probe_Set_v1.0_GRCh38-2020-A (10x Genomics). The rest of the analysis was performed using Seurat v4 [26]and standard R packages. Custom filtering at the cell level was performed with between 3,000 and 30,000 UMI counts and between 2,000 and 8,000 distinct features (genes), a maximum of 5% mitochondrial and ribosomal reads, as well as a doublet score (scrublet-0.2.3 [27]) below 0.15. The Seurat integration algorithm [28] was used to combine the six individual samples into one dataset. Cluster annotation was performed based on differentially expressed marker genes from published data [3, 29]. Single-sample Gene Set Enrichment Analysis (ssGSEA) was applied to snRNA-seq data using the escape R-package (doi:https://doi.org/10.18129/B9.bioc.escape). The msigdbr and getGeneSets functions were used to fetch and filter the corresponding pathway and ontology gene sets from the MSigDB. EnrichIt with default parameters was performed on the Seurat object, except for using 10,000 groups and a variable number of cores.

### Calcium live cell imaging

The fluorogenic calcium-binding dye Fluo-Forte (Enzo Life Sciences) is cell-permeable and, once inside the cells, is hydrolyzed to a cell membrane impermeable form. Increases in intracellular calcium led to higher fluorescence signals. Co-cultures were treated with Fluo-Forte according to the manufacturer’s protocols and stained with the fluorescently labeled antibody CD88-PE (1:200) for 1h at 37°C in PBS. Intracellular calcium levels were measured in real-time using a confocal microscope (Excitation: 488 nm, LSM 710 Zeiss, Jena, Germany). Cells were imaged for 20 seconds (1 image/second) to establish a baseline. After 20 images, cultures were treated with 0.1 mM ATP. The measurement was continued for an additional 40 frames to investigate the washout fluorescence intensity. Images were analyzed using ImageJ2 (Version 2.14.0/1.54f, NIH, Bethesda, USA). CD88 signals were used to determine microglial positions within the co-culture images. Each microglial cell was analyzed separately by calculating the pixel intensity of each frame normalized to the baseline level intensity (F/F0). Calcium responses were plotted as a function of time (frames), and a quantification of the maximal peak amplitude of calcium was performed upon application of ATP (frame 20).

### Statistics

Statistical analysis for single-nucleus RNA-seq data during ssGSEA (Figure 3) and to compare average fold-changes (Figure 2G) was performed by applying a double-sided Wilcoxon test (rstatix::wilcox_test) with default parameters to statistically compare scores between cells from Parkin-PD patients and controls in a gene-wise fashion (Table S4).

The remaining data were analyzed using GraphPad Prism (Version 8, GraphPad Software, La Jolla, USA). One-way ANOVA or Two-way ANOVA followed by multiple comparison tests were performed as indicated. The p-values are illustrated in figures as *p<0.05, **p<0.01, ***p<0.001, ****p<0.0001.

## RESULTS

### Generation of iPSC-derived DA neuron-microglia co-cultures

To identify gene expression variation underlying the pathological processes caused by the loss of Parkin, we employed iPSC lines derived from three PD patients carrying biallelic *PRKN* mutations (Parkin-PD-1; Parkin-PD-2; Parkin-PD-3) and three healthy controls (ctrl-1; ctrl-2; ctrl-3) (Table S1, Table S2, Figure S1). All iPSC lines were differentiated separately into precursors of DA neurons and microglia and matured under co-culturing conditions (Figure 1A) as previously described [20, 21, 23], with each DA neuron donor combined with the same microglia donor. Microglia precursor cultures were characterized by the expression of myeloid surface markers CD11b, CD14, CD45, and CD88 (Figure S2) and plated onto DA 50-day-old neuronal precursors. Co-cultures were analyzed after 14 days of combined maturation for the presence of key marker proteins. Immunofluorescence staining showed the expression of DA marker tyrosine hydroxylase (TH), neuronal class III beta-tubulin (TUJ1), and microglial ionized calcium-binding adaptor molecule 1 (IBA1) (Figure 1B). Western blotting confirmed the expression of TH and IBA1 as well as the neuronal markers microtubule-associated protein 2 (MAP2) and neuron-specific enolase (NSE) (Figure 1C). There was no significant difference between Parkin-PD and control lines when comparing protein levels of these key tissue markers in the cell cultures (Figure 1D).

**Figure 1.**
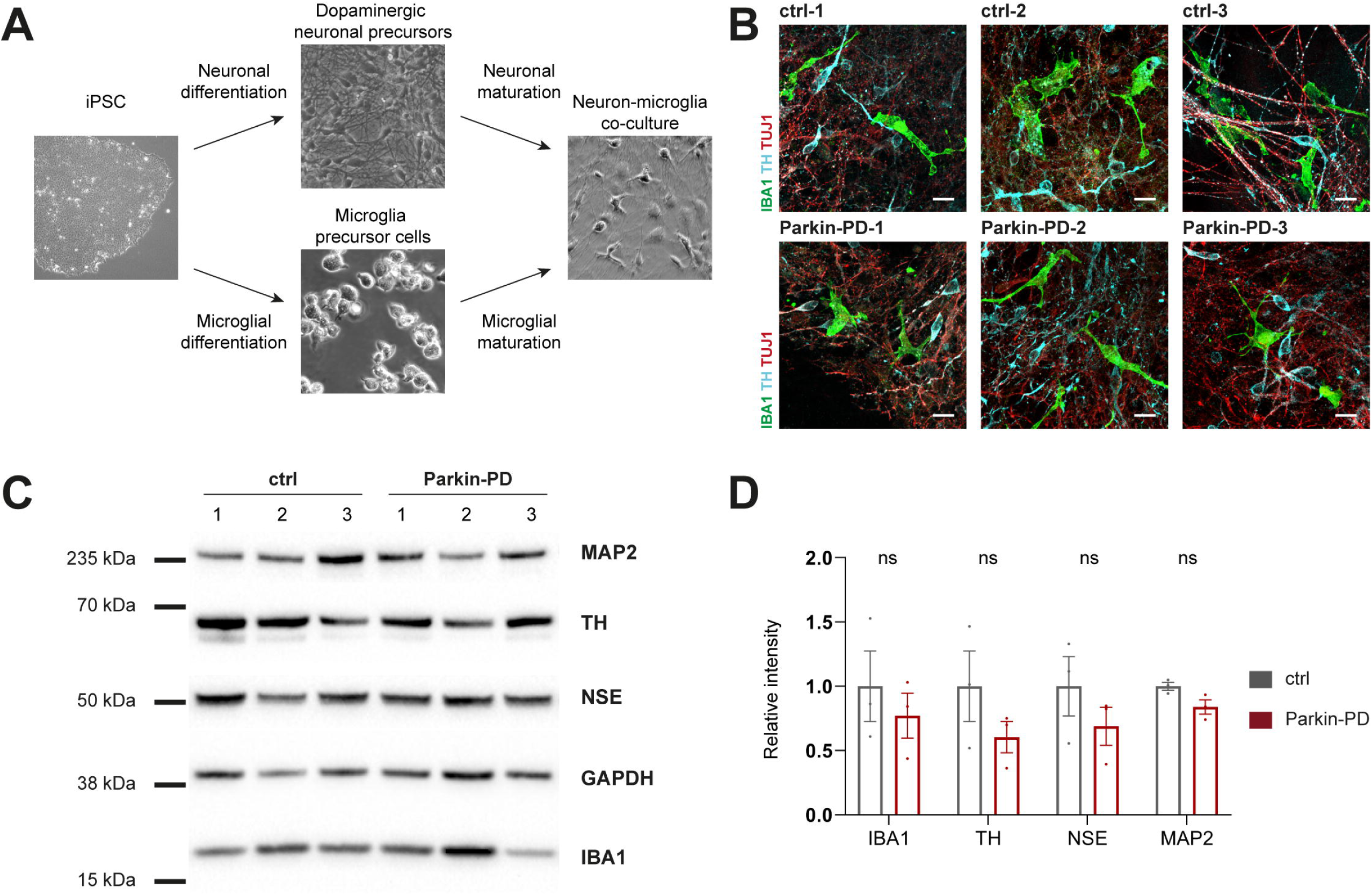
Characterization of iPSC-derived neuron-microglia co-cultures. Data are shown for differentiated cell cultures derived from three healthy control individuals (ctrl-1, ctrl-2, ctrl-3) and three Parkin-linked PD patients (Parkin-PD-1, Parkin-PD-2, Parkin-PD-3). **(A)** Differentiation scheme of neuron-microglia co-cultures. **(B)** Immunofluorescence staining for microglial ionized calcium-binding adaptor molecule 1 (IBA1, green), neuron-specific class III beta-tubulin (TUJ1, red), and the dopaminergic marker tyrosine hydroxylase (TH, blue). Images are representative of n=5 independent differentiation experiments. Scale bar, 25 µm. **(C)** Western blot analysis of whole-cell lysates from co-cultures for the neuronal marker proteins microtubule-associated protein 2 (MAP2) and neuron-specific enolase (NSE), microglial IBA1, dopaminergic TH, and GAPDH as loading control. Whole blots are shown in Figure S3. **(D)** Signal intensity analysis of data shown in (C) relative to GAPDH; two-way ANOVA followed by Sidak’s multiple comparisons test; n=3 differentiations of 3 ctrl lines and 3 Parkin-PD lines; ns, not significant. Mean ± SEM.

### Single-nucleus RNA-seq reveals distinct cell clusters

Using Chromium Next GEM Single Cell 3L v3.1 chemistry on a Chromium X device (10x Genomics), we performed single-nucleus RNA-seq from *PRKN*-mutant and control DA neuron-microglia co-cultures (Figure 2A). The dataset after filtering includes 22,451 and 22,466 FACS-sorted high-quality single-nuclei transcriptomes from Parkin-PD patients and controls, respectively, with a median UMI count per cell of 8,605 and a median gene count per cell of 4,417.

**Figure 2.**
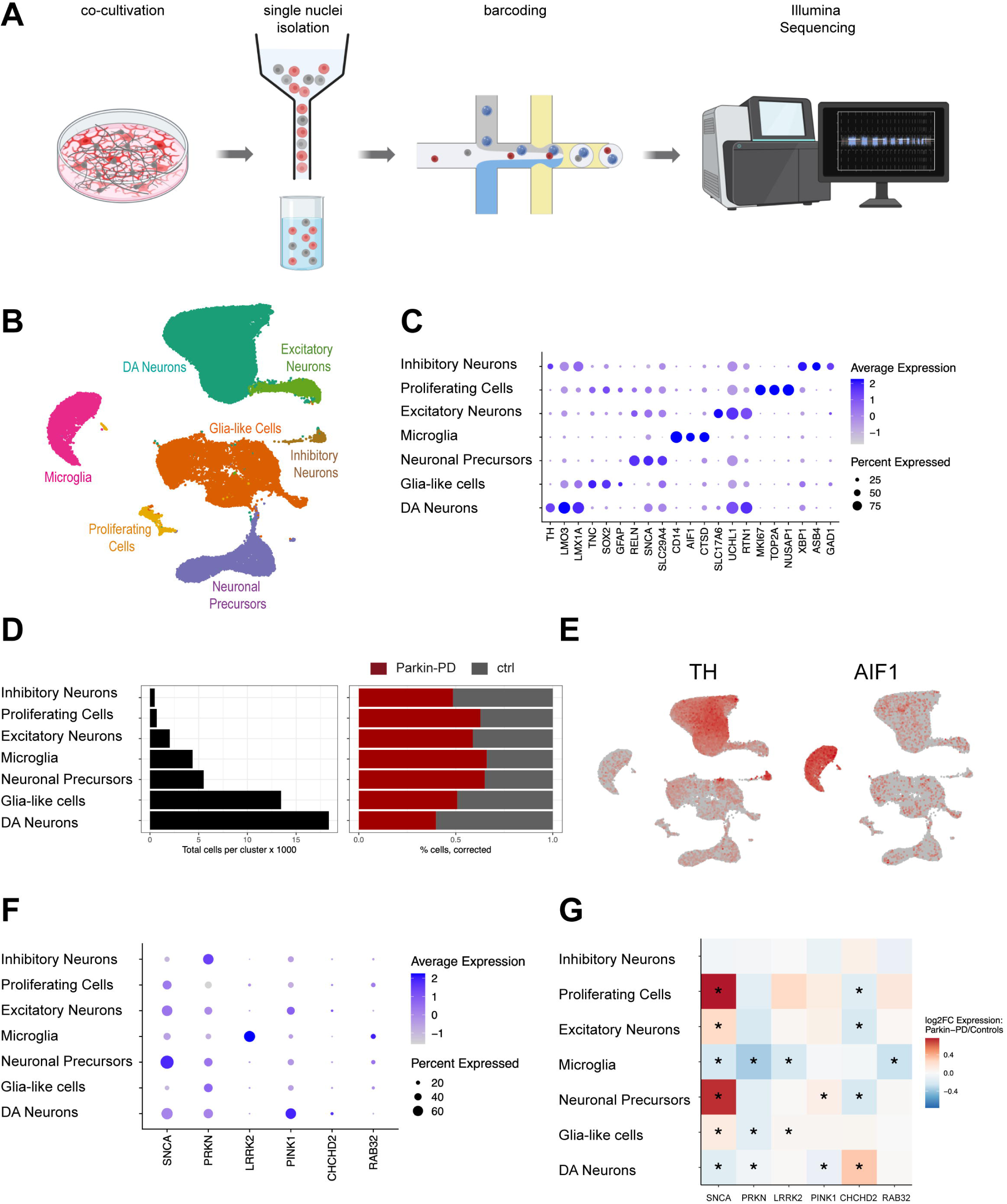
Cell type composition of neuron-microglia cultures. **(A)** Nuclei suspensions of co-cultures were sorted for intact nuclei using FACS, processed with the 10x Genomics platform for single nuclei barcoding, and sequenced with an Illumina sequencer. Created with BioRender.com. **(B)** UMAP embedding of 22,451 nuclei from three Parkin-PD patients and 22,466 nuclei from three healthy control individuals (ctrl). **(C)** Dot plot of cell type-specific representative marker genes. **(D)** Total number of profiled nuclei per cell type (left). The proportion of Parkin-PD and ctrl profiled cells per cell type (right). **(E)** Cell cluster expression for DA neuronal (*tyrosine hydroxylase*, *TH*) and microglial (*allograft inflammatory factor 1*, *AIF1*) markers. **(F)** Cell type-specific enrichment of genes associated with monogenic PD. **(G)** Cell type-specific average log2 fold change of expression in Parkin-PD vs ctrl samples is shown (Wilcoxon test, *FDR-corrected p-value<0.05).

We used the Seurat v4 internal, anchor-based integration approach to account for inter-individual variation during clustering and visualized the dataset in UMAP (Figure 2B). Louvain-based cluster identification revealed seven major cell types composed of neuronal, glial, and precursor cells. Overall, the cluster structure was driven by cell type identity, with cell type composition showing strong sample-specific variability. We used a panel of well-known marker genes from the literature [3, 29] to confirm the cell type identity of the clusters (Figure 2C). DA neurons were characterized by the expression of *TH*, *LMO3* and *LMX1A*. In addition, we identified other neuronal subtypes, inhibiting neurons expressing *XBP1*, *ASB4* and *GAD1* and excitatory neurons expressing *SLC17A6*, *UCHL1* and *RTN1*. Microglia clearly separated from the other clusters and expressed *CD14*, *AIF1* (encoding IBA1) and *CTSD*. Expression of *TNC*, *SOX2* and *GFAP* was characteristic of glia-like cells. Finally, we identified precursor cells that were clustered into neuronal precursors (*RELN*, *SNCA* and *SLC29A4*) and highly proliferating cells (*MKI67*, *TOP2A* and *NUSAP1*). All these cell types were detectable in both control and Parkin-PD cultures (Figure 2D). We found that neurons comprised 46.5% of total cells, with DA neurons accounting for 41.1% and inhibitory and excitatory neurons accounting for 1.1% and 4.4% of the total cell population, respectively. Further, 9.5% of the sampled cells were identified as microglia, 30.2% as glia-like cells, and the remaining cell types were composed of 12.0% neuronal precursors and 1.7% proliferating cells.

Our study focuses primarily on the cell clusters of DA neurons, the main affected neuronal subtype in PD, and microglia, the brain-resident immune cells (Figure 2E). We first explored the cell type specificity of a set of genes unequivocally linked to monogenic PD [30]. *SNCA, PRKN*, and *PINK1* show higher expression in DA neurons or neuronal precursors as compared to microglia, whereas *LRRK2* expression is specific to microglia (Figure 2F). In contrast, *CHCHD2* and *RAB32* did not show cell-type-specific enrichment. Next, we compared gene expression levels in Parkin-PD vs. control cell lines (Figure 2G). In DA neurons, *CHCHD2* was upregulated, whereas *PRKN*, *PINK1*, and *SNCA* were downregulated. In microglia, *LRRK2*, *PRKN*, *RAB32*, and *SNCA* were downregulated. Of note, the PD-linked genes *VPS35* and *PARK7* were not detectable at sufficient levels in the data set for further analysis. In the following, we examined the effect of Parkin deficiency on global gene expression in our co-culture model.

### Parkin deficiency affects Parkinson-related pathways in a cell type-specific manner

To gain cellular mechanistic insight, we evaluated biological pathways and relevant gene sets from the molecular signatures database (MSigDB) using single-sample gene set enrichment analysis (ssGSEA) [31–33], focusing on terms related to “Parkin”, “mitochondria”, “dopamine”, “inflammation”, and “calcium”. Among the key hits of 108 analyzed gene sets, we identified numerous cell type-specific dysregulations comparing Parkin-PD and controls (Figure S4, Table S4). A subset is shown exemplarily in more detail in Figure 3. In the categories “Parkin-associated” and “mitochondria-associated”, gene sets of the BioCarta “Parkin pathway” and the gene ontology biological pathway (GOBP) “Parkin-mediated stimulation of mitophagy in response to mitochondrial depolarization” showed dysregulation mainly in DA neurons, whereby Parkin-PD lines displayed decreased expression compared to controls (Figure 3A) [34–37].

**Figure 3.**
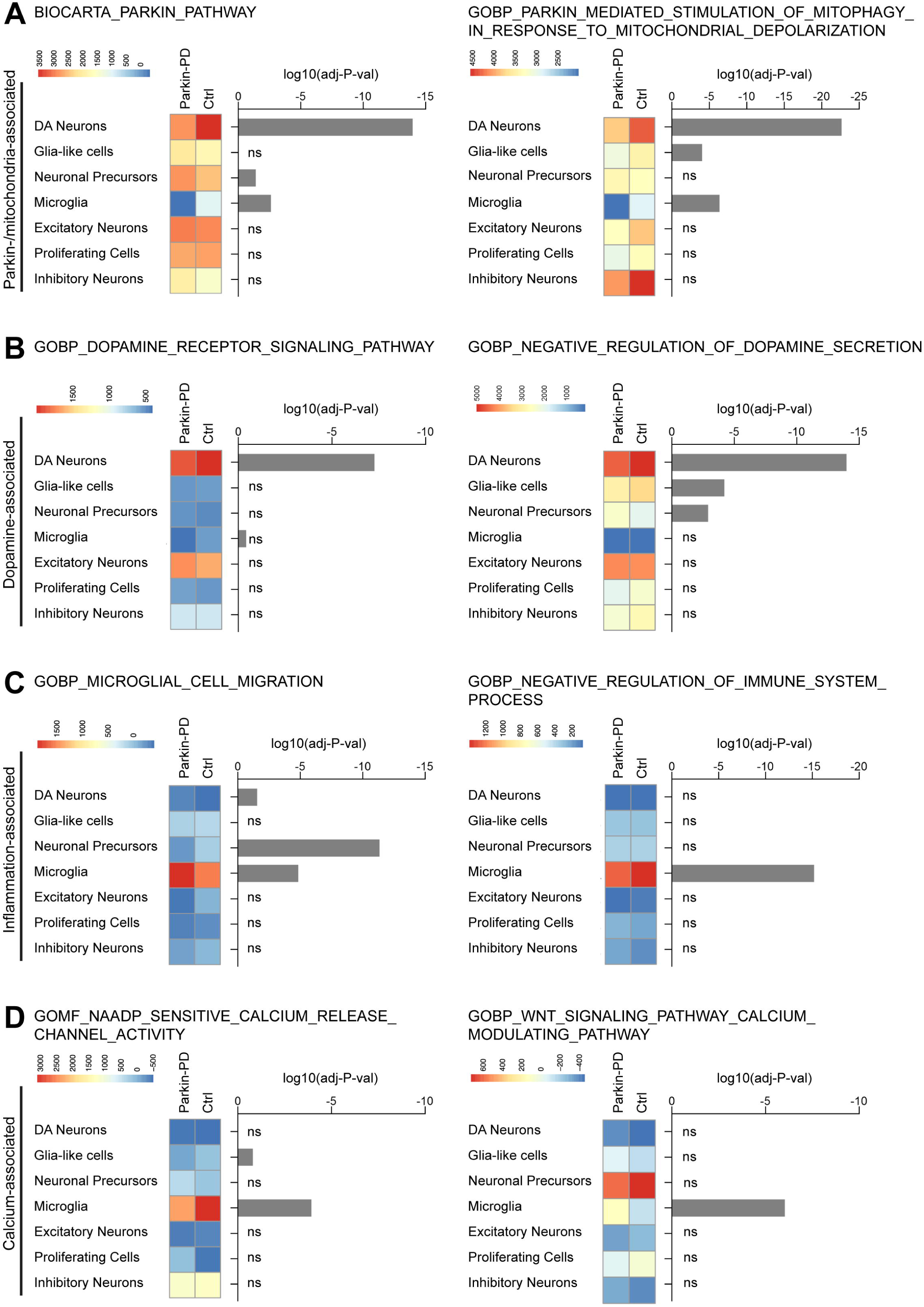
Differentially regulated biological pathways in neuron-microglia co-cultures. Heatmaps of biological pathways in the categories **(A)** Parkin-/mitochondria-associated; **(B)** Dopamine-associated; **(C)** Inflammation-associated; **(D)** Calcium-associated. P-values were determined by double-sided Wilcoxon test with FDR correction (Figure S4, Table S3).

In the category “dopamine-associated”, GOBP “dopamine receptor signaling pathway” and GOBP “negative regulation of dopamine secretion” gene sets displayed the highest enrichment in DA neurons compared to other cell types, supporting the DA neuron identity within our cell type calling (Figure 3B). Importantly, elevated enrichment was detected in control lines compared to Parkin-PD lines.c

In the category “inflammation-associated”, GOBP “microglia cell migration” and GOBP “negative regulation of immune system process” gene sets show strong enrichment in microglia compared to other cell types and significant differential enrichment in control compared to Parkin-PD lines (Figure 3C). This supports the microglial identity in our cell type calling and links Parkin deficiency to inflammatory signaling.

In the category “calcium-associated”, enrichment scores in gene ontology molecular function (GOMF) “NAADP sensitive calcium release channel activity” and GOBP “WNT signaling pathway calcium modulating pathway” gene sets suggest significant dysregulation in *PRKN*-mutant microglia (Figure 3D).

While iPSC-derived DA neurons harboring *PRKN* mutations have been previously studied, demonstrating impaired dopamine homeostasis and mitophagy defects [13–16, 38], the impact of *PRKN* mutations on human microglia remains entirely unknown. We therefore focused on microglial dysfunction and asked whether the transcriptomic disturbances also translate into cellular detectable phenotypes.

### *PRKN*-mutant microglia show enhanced MCP-1 secretion and altered IL-8 response

Based on observed dysregulation in microglial inflammation-associated pathways, we examined the release of cytokines in the co-culture medium from Parkin-PD patients and controls.

Under basal conditions, most cytokines were detected at low levels in undiluted cell culture medium samples in a multiplex assay (Figure 4A). Interestingly, levels of the monocyte chemotactic protein 1 (MCP-1, CCL2) were significantly increased in Parkin-PD compared to control samples. This observation is consistent with the upregulation of the GOBP “microglia cell migration” gene set in Parkin-PD lines, as indicated by elevated enrichment scores (Figure 3C). The gene set includes genes associated with chemotactic signaling (*CSF1, CX3CR1, STAP1, P2RX4, TREM2, CCL3, CX3CL1, P2RY12*).

**Figure 4.**
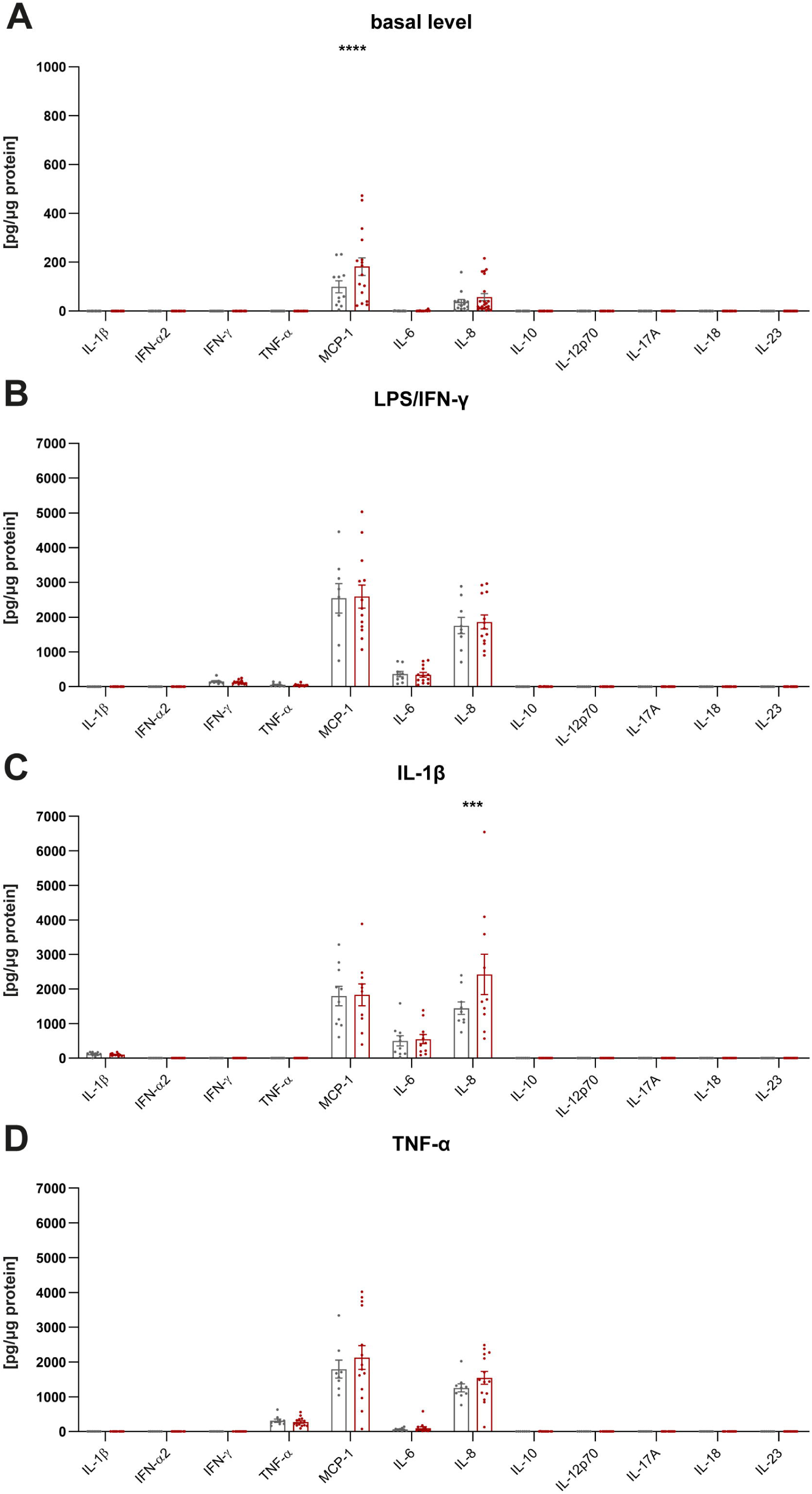
Cytokine profiling of neuron-microglia co-cultures. Simultaneous measurement of 12 cytokines in cell culture media by FACS using Legendplex Human Inflammation Panel 1 under **(A)** basal conditions and after treatment with **(B)** LPS/IFN-γ, **(C)** IL-1β, and **(D)** TNF-α. Cell culture media of three *PRKN*-mutant lines (n=22 samples, red) and three healthy control lines (n=15 samples, grey) were analyzed (n=10 differentiations). Asterisks indicate statistically significant differences between cultures using a two-way ANOVA followed by Sidak’s multiple comparisons test (*** p<0.001, ****p<0.0001). Mean ± SEM.

To investigate the co-cultures upon microglial activation, they were treated with inflammatory stressors (Figure 4B-D). IFN-γ was used together with LPS to mimic a microglial activation state similar to an *in vivo* situation in which IFN-γ acts in the co-presence of damage-associated molecular patterns. This resulted in a strong response, and cells secreted the cytokines MCP-1, IL-6, and IL-8 at high levels into the culture medium, but without differences between mutant and control cultures. Further, IL-1β or TNF-α were used to stimulate the co-cultures. Polymorphisms in the *IL1B* and *TNF* genes have been identified as risk factors of PD [5], and both cytokines have been suggested as important mediators of the functional effects of neuroinflammation on neurodegeneration in PD models [39]. When comparing Parkin-PD lines with controls, the only statistically significant difference in cytokine production was observed for the chemokine IL-8, which increased in Parkin-PD samples upon IL-1β treatment (Figure 4C).

### *PRKN*-mutant microglia reveal impaired calcium release

Next, we explored variations of calcium-associated pathways specific to *PRKN*-mutant microglia by measuring intracellular calcium levels in living cells. Co-cultures were treated with the fluorogenic calcium-binding dye Fluo-Forte and were co-stained for the microglial cell surface marker CD88 (C5a anaphylatoxin chemotactic receptor 1, C5aR1). Confocal microscopy revealed lower intracellular calcium levels in Parkin-PD microglia compared to controls under basal conditions (Figure 5). An important modulator of microglial functions, including proliferation, migration, phagocytosis, and secretion of inflammatory mediators, is extracellular ATP, which increases intracellular free calcium concentration [40]. Notably, we found markedly reduced intracellular calcium release in ATP-stimulated *PRKN*-mutant microglia compared to controls (Figure 5), suggesting impaired calcium signaling due to loss of Parkin.

**Figure 5.**
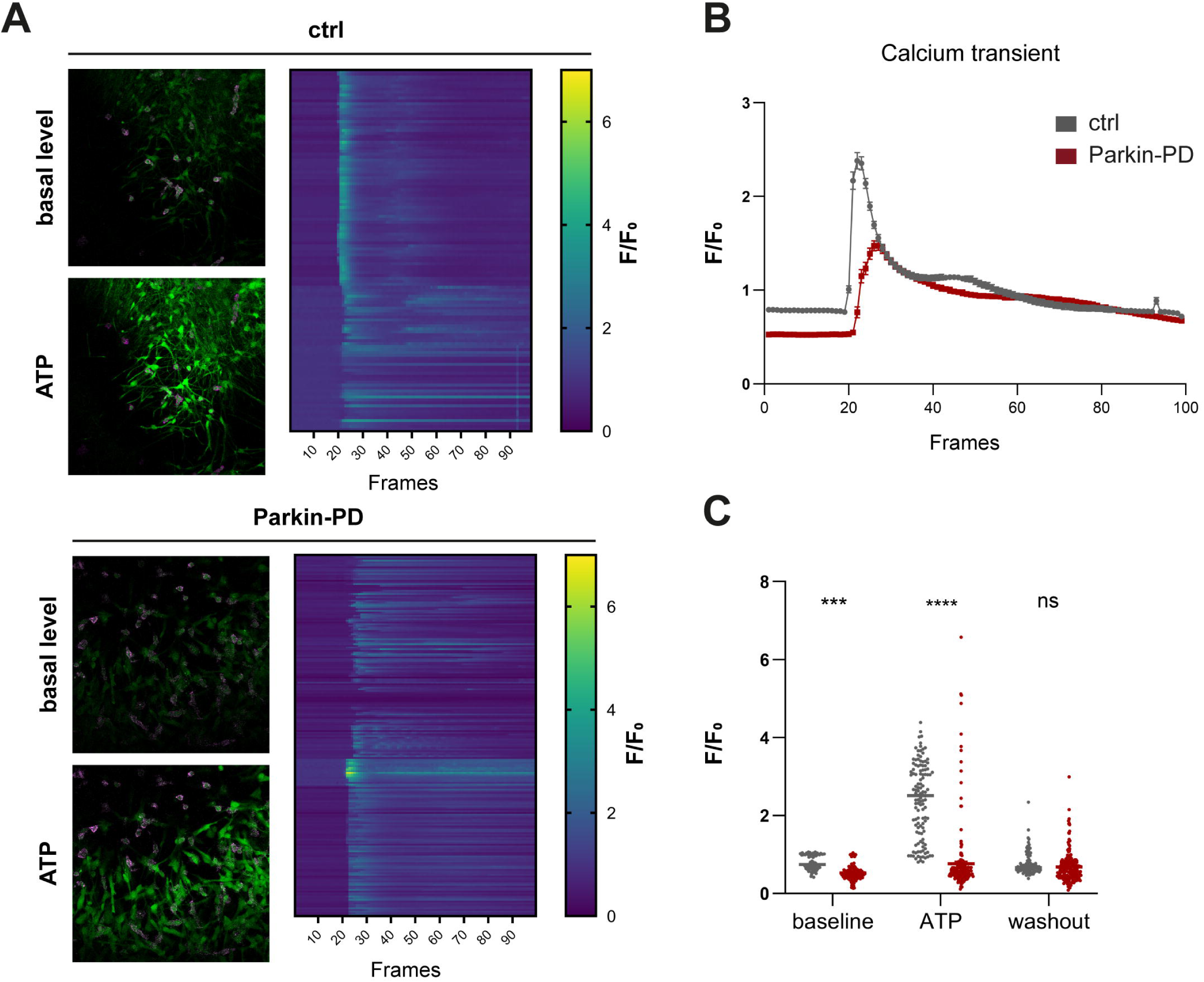
Calcium imaging of neuron-microglia co-cultures. **(A)** Measurement of calcium release in neuron-microglia co-cultures. Representative images with calcium fluorophore (green) under baseline conditions and after treatment with 0.1 mM ATP. Microglia were co-stained with CD88-PE (purple). The heatmaps show calcium responses in control (ctrl) and Parkin-PD lines over time. **(B)** Examples of evoked responses (ΔF/F0) from ctrl (grey) and Parkin-PD (red) microglia were plotted as a function of time (frames). **(C)** Quantification of the calcium measurements is shown: Ctrl (grey) n=122 cells (ctrl-1; ctrl-2), 3 independent measurements; Parkin-PD (red) n=217 cells (Parkin-PD-1; Parkin-PD-3), 5 independent measurements. Asterisks indicate significant differences between ctrls and Parkin-PD cultures analyzed by two-way ANOVA followed by Sidak’s multiple comparisons test (*** p<0.001, **** p<0.0001, not significant (ns)). Mean ± SEM.

## DISCUSSION

In this study, we generated single-nucleus transcriptomes from iPSC-derived *PRKN*-mutant and healthy control neuron-microglia co-cultures. Pathway analysis revealed cell type-specific dysregulated biological processes, including *PRKN*-containing gene sets, as well as gene expression variation that controls cellular dopamine homeostasis, calcium homeostasis, and inflammatory signaling. Our subsequent functional analysis showed elevated chemokine levels and impaired calcium homeostasis in *PRKN*-mutant co-cultures. To date, only a limited number of studies have reported single-cell RNA-seq data for iPSC-derived models of genetic PD. In *GBA1*-linked PD, single-cell sequencing identified HDAC4 as a regulator of PD-related cellular phenotypes in DA neurons [41]. Upregulation of glycolysis and ubiquitin-proteasomal degradation genes was detected in *SNCA*-mutant DA neurons under stress compared to controls [19]. Single-cell transcriptomics of *PINK1*-mutant DA neurons revealed a core molecular network of PD that encompasses PD-associated pathways [42]. In our study, we increased the model’s complexity and examined *PRKN*-mutant DA neurons in co-culture with microglia. Importantly, our nuclei assignment was consistent with the pathway analysis in which cell type-specific pathways, such as dopamine receptor signaling or microglial migration, were enriched in DA neurons or microglia, respectively. In a first expression analysis, we focused on genes implicated with high confidence in monogenic PD. *Coiled-coil-helix-coiled-coil-helix domain containing 2* (*CHCHD2*) has been linked to autosomal dominant PD and encodes a protein residing in the mitochondrial intermembrane space [43, 44]. Notably, we found *CHCHD2* upregulated in *PRKN*-mutant DA neurons. This is consistent with our recent findings in TH-positive sorted DA neurons, in which *CHCHD2* upregulation was identified as one of the most significant traits when comparing *PRKN*-mutant and control cells (unpublished observation).

Based on previous findings, we focused our pathway analysis on terms related to “Parkin”, “mitochondria”, “dopamine”, “inflammation”, and “calcium”. The results are derived from a limited number of samples, including three patients and three controls. Therefore, we used enrichment score ranking per cell when implementing ssGSEA across the entire dataset, rather than a more robust, less type I error-prone pseudo-bulk analysis on this small set of samples to identify significant differences.

In *PRKN*-mutant DA neurons, our dataset revealed dysregulated gene expression for mitochondria- and dopamine-linked pathways. Mitochondrial dysfunction is a major theme in PD [45] that unifies phenotypes caused by mutations in most PD-linked genes. Loss of Parkin in iPSC-derived neurons leads to reduced mitochondrial volume fraction, decreased complex I activity, altered mitochondrial network morphology, and deficits in the mitochondrial biogenesis pathway [9–11]. Impaired dopamine metabolism has also been directly linked to Parkin dysfunction [46]. In iPSC-derived DA neurons, loss of Parkin reduced dopamine uptake and increased spontaneous dopamine release [15]. Importantly, our study provided insights into two *PRKN*-containing gene sets (“Parkin pathway” and “Parkin mediated stimulation of mitophagy in response to mitochondrial depolarization”) that were dysregulated predominantly in *PRKN*-mutant DA neurons and not the other cell types. Parkin function appears to be specifically relevant to DA neurons in both pathways. As Parkin is ubiquitously expressed in most cells of the human body, it remains a challenge in PD research to identify Parkin functions that are selectively vital for DA neurons.

In *PRKN*-mutant microglia, we found changes in the expression levels of inflammation- and calcium-linked pathways. Few studies have investigated *PRKN*-mutant microglia to date [47]. Activation of the NRLP3 inflammasome was observed in parkin-null murine microglia cells exposed to LPS [48]. In our previous study, Parkin-deficient neuron-microglia co-cultures showed increased IL-6 gene expression upon exposure to mtDNA/LPS [11]. We now performed a cytokine panel under basal conditions and found increased MCP-1 levels in the cell culture medium of Parkin-PD co-cultures. MCP-1 is among the most highly expressed chemokines during inflammation and has been associated with many neurodegenerative disorders through the regulation of monocyte chemotaxis [49]. Notably, MCP-1 has been shown to correlate with the progression of idiopathic PD in CSF samples [50]. In addition, increased CSF levels of IL-6, TNF-α, IL-1β, CRP, and MCP-1 were revealed in patients with idiopathic PD compared to controls in a recent meta-analysis of inflammatory biomarkers [51].

Treatment with IL-1β resulted in increased IL-8 release in Parkin-PD cultures in our study. Notably, both IL-8 and MCP-1 function in the chemotaxis pathway. When neuronal damage is detected by microglia, ramified microglia become activated and migrate toward the site of pathology, using a chemoattractant gradient as a directional cue [52].

In terms of Parkin-related calcium dysregulation, we recently reported that calcium distribution at mitochondria-endoplasmic reticulum (ER) contact sites was disrupted in *PRKN*-mutant neurons [17]. Calcium also regulates several microglial processes, including proliferation, migration, phagocytosis, and secretion of inflammatory mediators. Extracellular ATP triggers these processes by activating specific receptors, thereby increasing intracellular free calcium concentration in human microglia [40]. Recently, calcium imaging studies demonstrated a reduction of ATP-induced ER calcium release in mouse parkin-deficient astrocytes [53]. We found decreased intracellular calcium levels in human *PRKN*-mutant microglia compared to controls. ATP treatment resulted in reduced intracellular calcium release, indicating disturbed calcium homeostasis. In contrast, parkin-knockout mouse embryonic fibroblasts showed increased ER calcium release upon ATP treatment compared to control cells [54]. Because calcium signaling is regulated in a cell type-specific manner, diverse effects may be observed across different cell types and/or species.

In summary, our unique dataset underscores the importance of cell type-specific analysis. We provide a human disease model that verified a broad range of Parkin-linked phenotypes in DA neurons and uncovered unknown microglial dysfunction. Further in-depth functional analysis will be needed to elucidate the combined impact of DA neuron and microglial impairment on the pathogenesis in *PRKN*-mutant brains.

## CONCLUSION

Profiling at the single-cell level resolved distinct cell subpopulations, enabling us to identify cell type-specific pathways and gene sets associated with PD. We validated our key findings by demonstrating increased chemokine release and disturbed intracellular calcium levels in DA neuron-microglia co-cultures harboring *PRKN* mutations. This unique dataset provides a basis for new research approaches to tease apart the cell type-specific impairments underlying PD.

## LIST OF ABBREVIATIONS

CD: Cluster of Differentiation
CHCHD2: Coiled-coil-helix-coiled-coil-helix domain containing 2
CSF: Cerebrospinal fluid
Ctrl: control
DA: dopaminergic
EB: Embryoid bodies
ER: Endoplasmic Reticulum
FACS: Fluorescence Activated Cell Sorting
GEM: Gel Beads-in-emulsion
GOBP: gene ontology biological pathway
GOMF: gene ontology molecular function
hPSCreg: Registry for human pluripotent stem cells
IFN-γ: Interferon gamma
IL-1β: Interleukin-1 beta
IL-8: Interleukin-8
iPSC: induced pluripotent stem cell
LPS: Lipopolysaccharide
MCP-1: Monocyte chemotactic protein 1
mtDNA: mitochondrial DNA
PE: Phycoerythrin
PD: Parkinson’s Disease
RNA-seq: RNA sequencing
RT: Room Temperature
SDS: Sodium Dodecyl Sulfate
TNF-α: Tumor necrosis factor alpha
TH: Tyrosine hydroxylase
UMAP: Uniform Manifold Approximation and Projection

## DECLARATIONS

### Ethics approval and consent to participate

This study was conducted in accordance with the Declaration of Helsinki and was approved by the Ethics Committee of the University of Luebeck (protocol code 16-039). Informed consent was obtained from all individuals involved in this study.

### Consent for publication

Not applicable

### Availability of data and materials

The datasets used and/or analyzed during the current study are available from the corresponding author on reasonable request.

## Authors’ contributions

Conceptualization, E.K., C.K. and P.S.; methodology, E.K., K.H., F.R., L.S.G., A.G., S.A.C. and P.S.; formal analysis, E.K., K.H. and P.S.; investigation, E.K., K.H., F.R. and P.S.; writing—original draft preparation, E.K., K.H. and P.S.; writing—review and editing, E.K., K.H., F.R., L.S.G., D.A.F, A.G., S.A.C., M.S., C.K. and P.S.; funding acquisition, C.K. and P.S. All authors read and approved the final manuscript.

## Acknowledgments

We thank Sven Geisler (Institute for Systemic Inflammation Research, University of Luebeck, Germany) and Victor Krajka (Institute of Microtechnology, Technical University Braunschweig, Germany) for experimental advice and technical assistance.

