## Supplementary Material for "Single-cell resolution uncovers cell type-specific dysregulation in Parkin-deficient neuron-microglia co-cultures"

### SUPPLEMENTARY INFORMATION

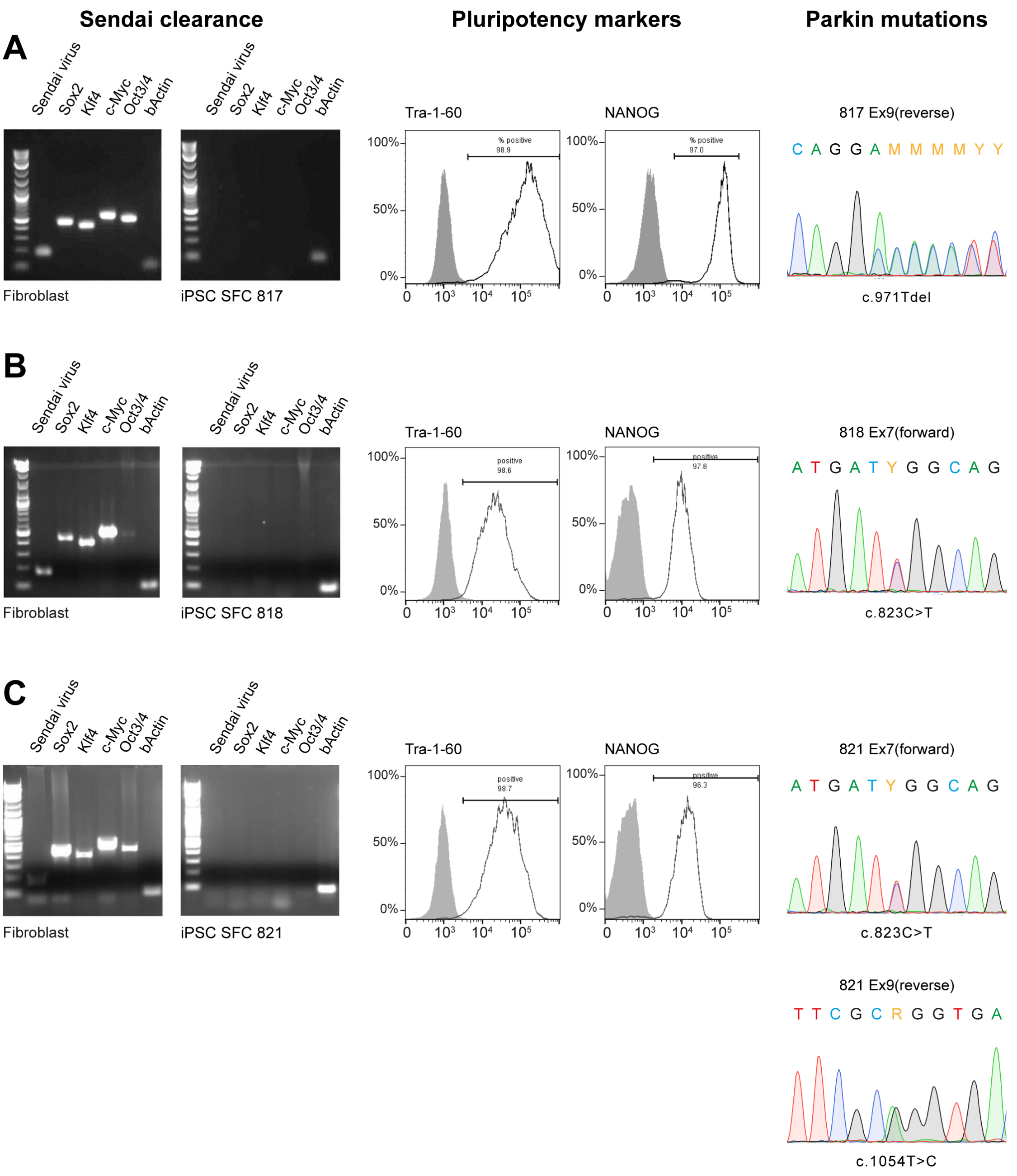


**Figure S1: Characterization of *PRKN*-mutant iPSC lines. (A)** iPSC SFC817, **(B)** iPSC SFC818, and **(C)** iPSC SFC821. Left panel: Gene expression analysis of the Sendai virus reprogramming factors (Sox2, Klf4, c-Myc, and Oct3/4) directly after transduction (fibroblast) and after ten passaging steps of isolated iPSC clones. Middle panel: Analysis of pluripotency markers (Tra-1-60, NANOG) in fibroblasts (grey) and iPSC lines (white) via FACS. Right panel: Sanger sequencing of iPSC lines to verify single nucleotide *PRKN* mutations. Whole exon deletions were detected by using Multiplex Ligation-dependent Probe Amplification (data not shown).

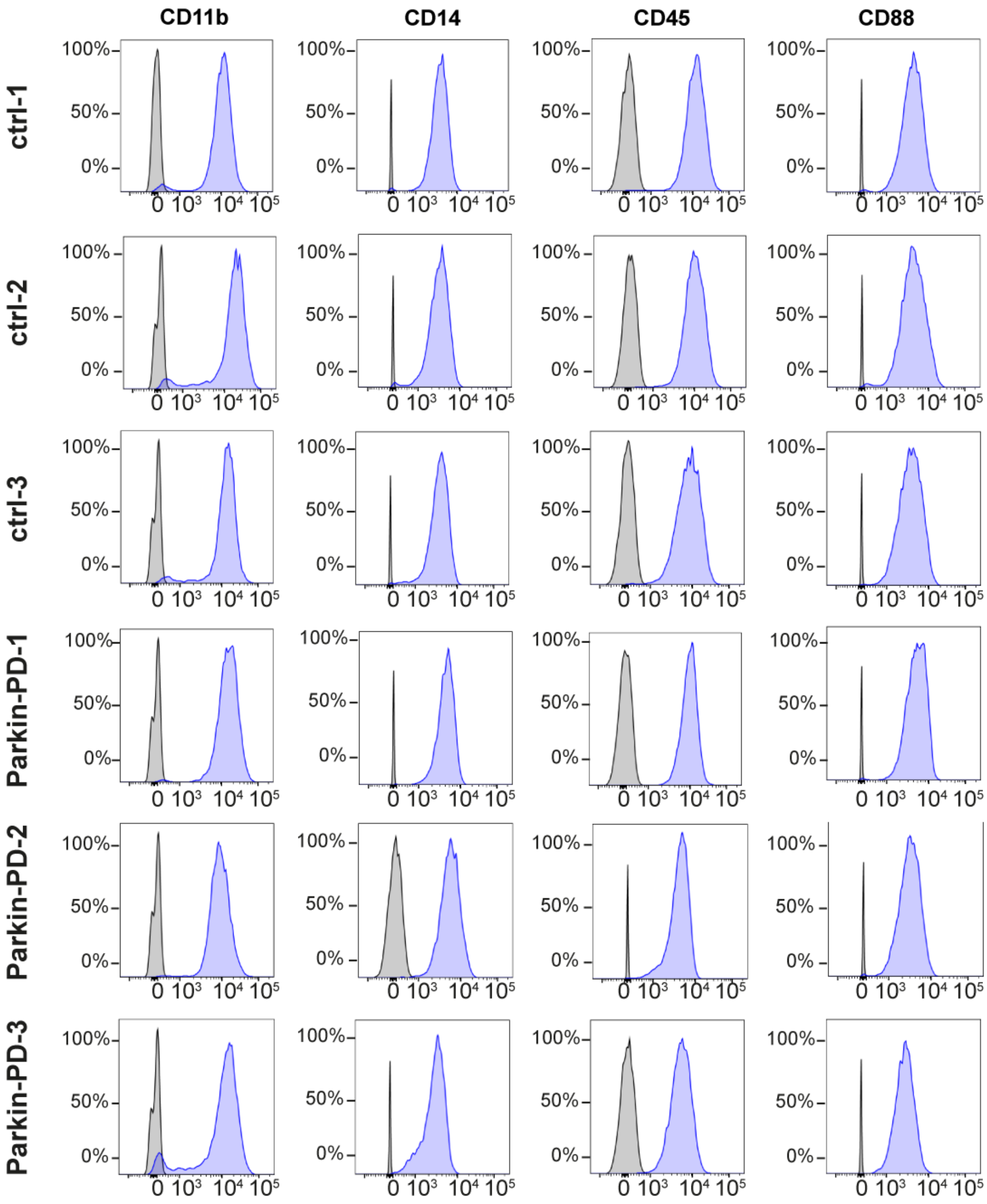


**Figure S2: Characterization of microglial progenitor cells.** Flow cytometry of iPSC-derived myeloid precursor cells (purple) and undifferentiated iPSCs (grey) for myeloid cell surface markers CD11b, CD14, CD45, and CD88. Graphs are representative of n=5 independent differentiation experiments.

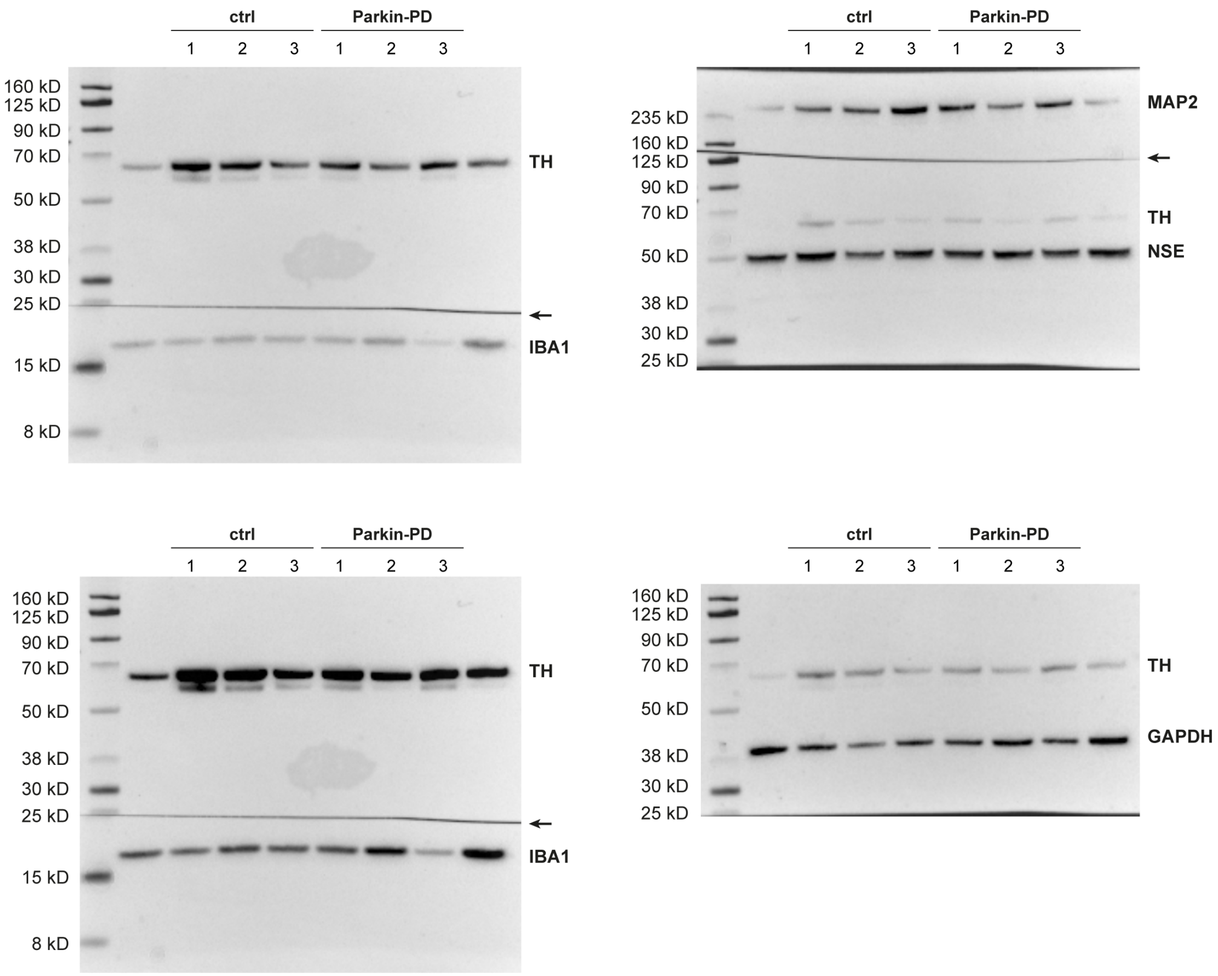


**Figure S3: Uncropped Western blots. (A)** Western blot analysis of microglial marker protein Ionized calcium-binding adaptor molecule 1 (IBA1) and neuronal marker proteins microtubule-associated protein 2 (MAP2), neuron-specific enolase (NSE), tyrosine hydroxylase (TH), and GAPDH. The membrane was cut as indicated by the arrows and the pieces were incubated separately with the antibodies specified. Exposure times: upper left panel 4 sec, lower left panel 30 sec, upper right panel 4 sec, lower right panel 1 sec.

**Figure S4:** Overview of biological pathways and relevant gene sets from the molecular signatures database (MSigDB) using single-sample gene set enrichment analysis (ssGSEA). Statistical analysis in Table S4.

**Table S1. iPSC control lines**

| **ID of iPSC lines** | SFC086-03-03 | SFC089-03-07 | SFC156-03-01 |
|---|---|---|---|
| | ctrl-1 | ctrl-2 | ctrl-3 |
| **Sex** | female | female | male |
| **Age at biopsy** | 57 | 64 | 75 |
| **Reprogramming method** | Sendai virus | Sendai virus | Sendai virus |
| **iPSC clone characterization** | https://hpscreg.eu/cell-line/STBCi052-C | https://hpscreg.eu/cell-line/STBCi053-A | https://hpscreg.eu/cell-line/STBCi101-A |

**Table S2. iPSC *PRKN*-mutant lines**

| **ID of iPSC lines** | SFC817-03-06 | SFC818-03-04 | SFC821-03-01 |
|---|---|---|---|
| | Parkin-PD-1 | Parkin-PD-2 | Parkin-PD-3 |
| **Sex** | male | male | female |
| **Age at biopsy** | 75 | 57 | 35 |
| **Age at onset** | 64 | 15 | n/a |
| ***PRKN* mutations** | c.971Tdel; delEx7 | delEx4; c.823C>T | c.823C>T; c.1054T>C |
| **Zygosity** | compound heterozygous | compound heterozygous | compound heterozygous |
| **Clinical status** | PD | PD | PD |
| **Reprogramming method** | Sendai virus | Sendai virus | Sendai virus |
| **iPSC clone characterization** | in this study* | in this study* | in this study* |

* Figure S1

**Table S3. Antibodies**

| **Primary antibody** | **Cat. No.** | **Dilution** | **Manufacturer** |
|---|---|---|---|
| TH | AB152 | 1:2000 | Merck Millipore, Burlington (USA) |
| GAPDH | 2118S | 1:30,000 | Cell Signaling, Danvers (USA) |
| IBA1 | Ab178846 | 1:1000 | Abcam, Cambridge (USA) |
| TH | NB 300-110 | 1:500 | Novus Biologicals, Centennial (USA) |
| MAP2 | MAB3418 | 1:1000 | Merck Millipore, Burlington (USA) |
| NSE | Ab180943 | 1:30,000 | Abcam, Cambridge (UK) |
| CD45-APC | 21810456 | 1:100 | ImmunoTools, Friesoythe (Germany) |

| | | | |
|---|---|---|---|
| CD14-PE | 21620144 | 1:100 | ImmunoTools, Friesoythe (Germany) |
| CD88-PE | 344304 | 1:100 | Biolegend, San Diego (USA) |
| CD11b-APC | 301309 | 1:100 | Biolegend, San Diego (USA) |
