## Supplementary figures and images for "Single-cell resolution uncovers cell type-specific dysregulation in Parkin-deficient neuron-microglia co-cultures"

### Supplementary Figure S4

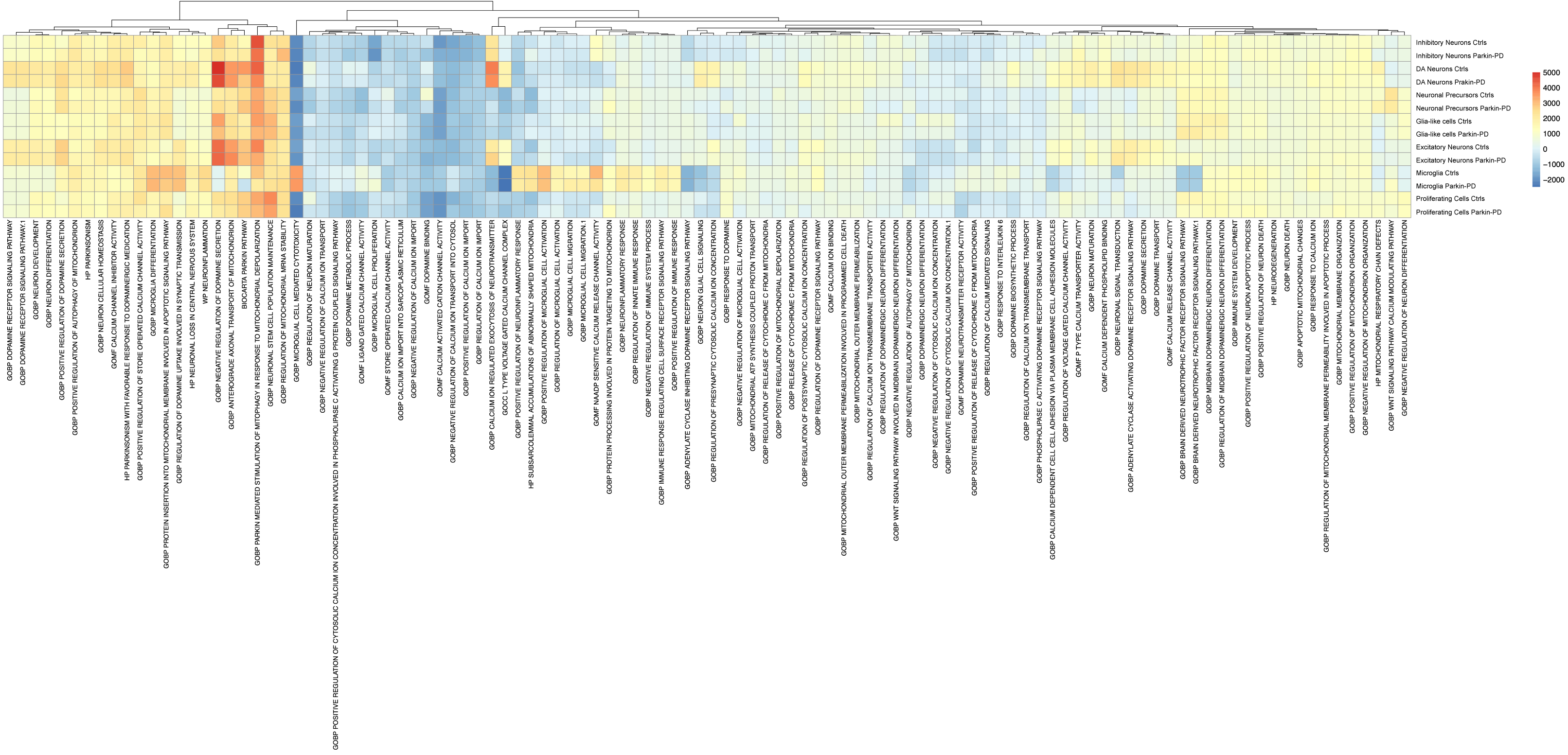
